# RegSNPs-Intron: A computational framework for prioritizing Intronic Single Nucleotide Variants in Human Genetic Disease

**DOI:** 10.1101/515171

**Authors:** Hai Lin, Katherine A. Hargreaves, Rudong Li, Jill L. Reiter, Matthew Mort, David N. Cooper, Yaoqi Zhou, Michael T. Eadon, M. Eileen Dolan, Joseph Ipe, Todd Skaar, Yunlong Liu

## Abstract

A large number of single nucleotide variants (SNVs) in the human genome are known to be responsible for inherited disease. An even larger number of SNVs, particularly those located in introns, have yet to be investigated for their pathogenic potential. Using known pathogenic and neutral intronic SNVs (iSNVs), we developed the regSNPs-intron algorithm based on a random forest classifier that integrates RNA splicing, protein structure and evolutionary conservation features. regSNPs-intron showed high accuracy in computing disease-causing probabilities of iSNVs. Using a high-throughput functional reporter assay called ASSET-seq (ASsay for Splicing using ExonTrap and sequencing), we validated regSNPs-intron predictions by measuring the impact of iSNVs on splicing outcome. Together, regSNPs-intron and ASSET-seq enable effective prioritization of iSNVs for disease pathogenesis. regSNPs-intron is available at https://regsnps-intron.ccbb.iupui.edu.

## Introduction

Prior to the advent of genome-wide association studies (GWAS), exploration of the relationship of genetic variants to human disease largely focused on non-synonymous single-nucleotide variants (SNVs) located in protein-coding regions. However, with the rise of next-generation sequencing, it is now clear that approximately one hundred times more variants are found in introns as compared to exons [1]. Importantly, intronic SNVs (iSNVs) can contribute to disease pathology by dysregulation of mRNA splicing [2-5]. For instance, over 3,000 disease-causing iSNVs in the Human Gene Mutation Database (HGMD) have been documented to impact splicing [6]. Most of the pathogenic variants locate close to the junction boundaries.

Intronic SNVs can impact alternative splicing by interfering with splice site recognition. For example, an intronic mutation near the 5’-splice site of exon 20 in the *IKBKAP* gene causes skipping of exon 20, resulting in malfunction of IKBKAP in 99.5% of familial dysautonomia (FD) cases [5, 7, 8]. Likewise, the intron 4 splice-donor site variant in the *adenomatous polyposis coli* (*APC*) gene causes skipping of exon 4, which can lead to colon cancer [9, 10]. Additionally, iSNVs may alter the binding affinities of RNA-binding proteins (RBP) to cis-regulatory elements [2, 11, 12]. For example, a G to A substitution within an intronic-splicing enhancer downstream of exon 3 in the growth hormone (GH1) gene can cause familial isolated GH deficiency type II (IGHD II) by suppressing the binding of splicing factors [13-15]. In addition, a recent support suggests that mis-splicing of the GALNS gene resulting from deep intronic mutations as a cause of Morquio a disease [16]. While iSNVs are densely distributed in the genome, only a limited proportion have been investigated for associations with altered biological functions [2].

Owing to the large number of intronic variants detected in next-generation sequencing and their complex RNA-splicing regulatory mechanisms, efficient bioinformatics algorithms are required to predict the potential impact of iSNVs and prioritize them for functional studies. One algorithm, SPANR (Splicing-based Analysis of Variants), was designed to evaluate how individual SNVs impact splicing regulation by predicting the maximum change (delta) in the percentage of inclusion (dPSI) of nearby exons induced by the SNVs. It extracts 1393 genomic features around the SNVs and predicts potential splicing outcomes by training a neural network with RNA-seq data from 16 human tissues [17]. However, SPANR was not designed to assess whether iSNVs result in deleterious phenotypes since it does not evaluate the impact of the resultant splicing change on protein function. Another widely used tool, CADD (Combined Annotation Dependent Depletion), predicts pathogenic variants with support vector machine (SVM) by combining annotations from multiple sources including: conservation scores, such as PhyloP; regulatory information, such as transcription factor binding; and protein-level predictions, such as SIFT and PolyPhen [18]. However, the annotations employed by CADD do not include RNA splicing data and provide limited information on protein function, as SIFT and PolyPhen only predict the consequences of SNVs based on amino acid substitutions. To this end, our earlier studies on small insertions/deletions (INDELs) [19, 20], alternatively spliced exons [21], and synonymous SNVs [22] [23], have indicated that simply substituting a stretch of amino acid residues does not necessarily imply altered protein function. Others have also reported that nucleotide substitutions can be bystander events, i.e. non-consequential to the phenotype [24]. Therefore, in order to consider the molecular implications of functional iSNVs, it is critical to integrate features that more accurately predict the effects of alternative-splicing events on protein structure and function.

In this study, we considered the impact of iSNVs on splicing regulation together with the variant-induced impact of alternatively-spliced exons on protein-structure features. We extracted pathogenic iSNVs from the HGMD and randomly selected neutral iSNVs from the 1000 Genomes Project [6, 25]. Using these data, we developed an algorithm based on random forest classifier to compute the disease-causing probabilities of iSNVs [26], which we have termed regSNPs-intron. This algorithm was also tested on an independent dataset selected from the ClinVar database [27]. In addition, we designed a high-throughput functional reporter assay, ASSET-seq (<u>AS</u>say for <u>S</u>plicing using <u>E</u>xon<u>T</u>rap and sequencing), to experimentally validate the effects of the predicted iSNVs on splicing regulation. regSNPs-intron and ASSET-seq as presented here can be used in tandem to prioritize and screen potential pathogenic iSNVs to better understand their roles in complex disease.

## Results

### Datasets

To compute the pathogenic probabilities of iSNVs, we constructed a training set by combining the manually-curated pathogenic iSNVs in the Human Gene Mutation Database (HGMD) and neutral iSNVs from the 1000 Genomes Project (Figure 1 and Figure S1). Neutral iSNVs documented in the 1000 Genomes dataset were derived from genome-sequencing data from 2,500 individuals lacking obvious clinical phenotypes [1]. In order to minimize the false negatives in our training set, we only selected those iSNVs with minor allele frequency (MAF) greater than 10%. This selection resulted in 2,438 pathogenic and 2,104,613 neutral iSNVs.

**Figure 1.**
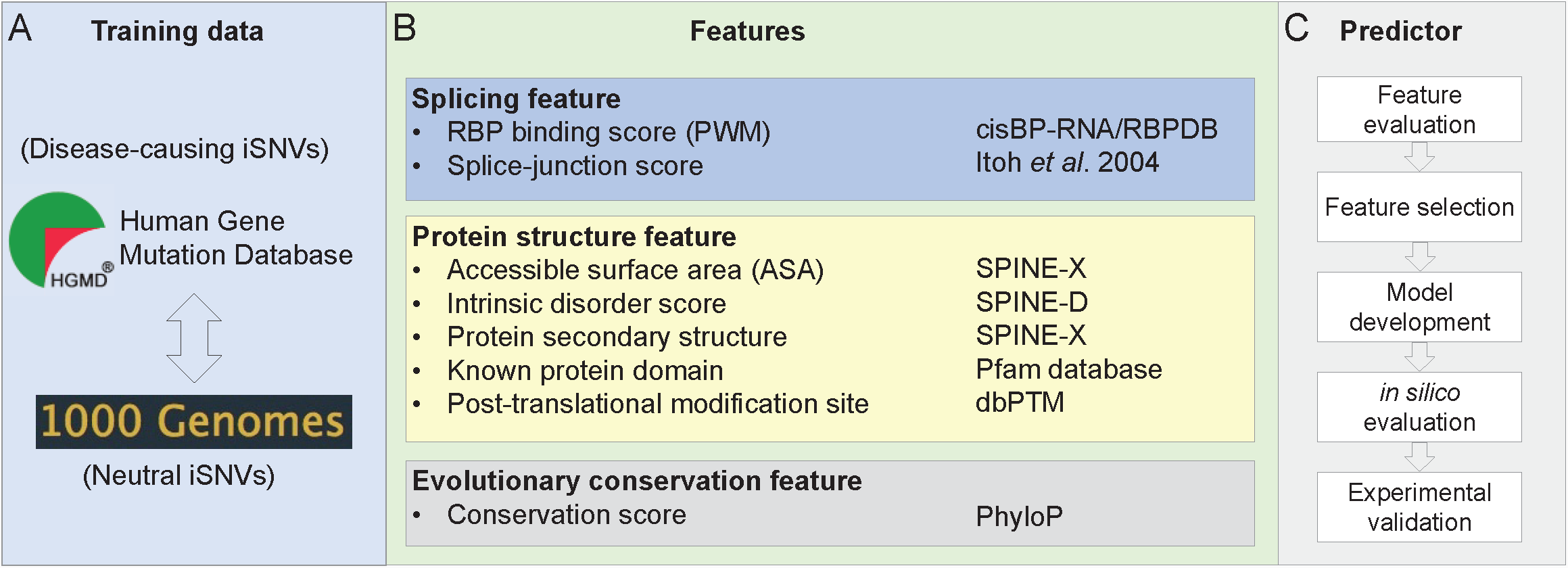
Workflow. (**A**) Data sources for model building. Pathogenic and neutral iSNVs were collected from HGMD and 1000 Genomes Project data, respectively, for model-training. (**B**) Three categories of features, namely RNA splicing, protein structure, and evolutionary conservation, were considered for the iSNVs. Data sources for each of these features are listed on the side. (**C**) Logic flow for prediction model development, as well as model evaluation and result validation.

Since the proximity of an iSNV to the splice-junction site (ss) can impact the splicing outcome via different molecular mechanisms, we further divided the selected iSNVs into those proximal and distal to the splice-junction sites. For example, variants proximal to the splice-junction sites (on-ss) may directly interfere with spliceosome formation, while iSNVs distal from the junction sites (off-ss) may affect the binding of regulatory RNA-binding proteins (RBPs). Splice-junction sites were defined as the upstream 13-bp for 3′-acceptor sites and downstream 7-bp for 5′-donor sites [28]. In total, there were 1,865 on-ss and 573 off-ss pathogenic iSNVs and 3,386 on-ss and 2,104,613 off-ss neutral iSNVs. Our data show that pathogenic variants were found more frequently in regions proximal to splice-junction sites compared to neutral variants (Figure S2). To avoid potential bias introduced by the variable distances of the proximal iSNVs from junction sites, we randomly selected 852 off-ss neutral iSNVs from the 1000 Genomes dataset by matching the distance distribution of the pathogenic HGMD variants. This ensured more balanced datasets with similar distance distributions between the pathogenic and neutral variants.

For each of the pathogenic and neutral on-ss and off-ss datasets, we randomly selected two-thirds of the data for use as the training set to build a random forest classifier (Figure S1). The remaining one-third of the data were used as the test set. To further test the model performance, we also extracted pathogenic and neutral iSNVs from the ClinVar database as an independent test set [27], which included 121 on-ss and 51 off- ss pathogenic iSNVs and 167 on-ss and 883 off-ss neutral iSNVs (see Methods section for details).

In order to select features that optimally discriminate pathogenic and neutral iSNVs, we classified all features into three categories: (i) splicing features, characterizing how individual iSNVs affect splicing regulation; (ii) structural features, evaluating how the iSNV-induced inclusion/exclusion of alternatively-spliced exons affect protein functions; and (iii) evolutionary-conservation features, nucleotide base-wise conservation scores of 99 vertebrate genomes (Figure 1). Most of the features showed significant power in separating the pathogenic and neutral iSNVs, based on the Wilcoxon rank-sum test (Figure 2 and Table S1).

**Figure 2.**
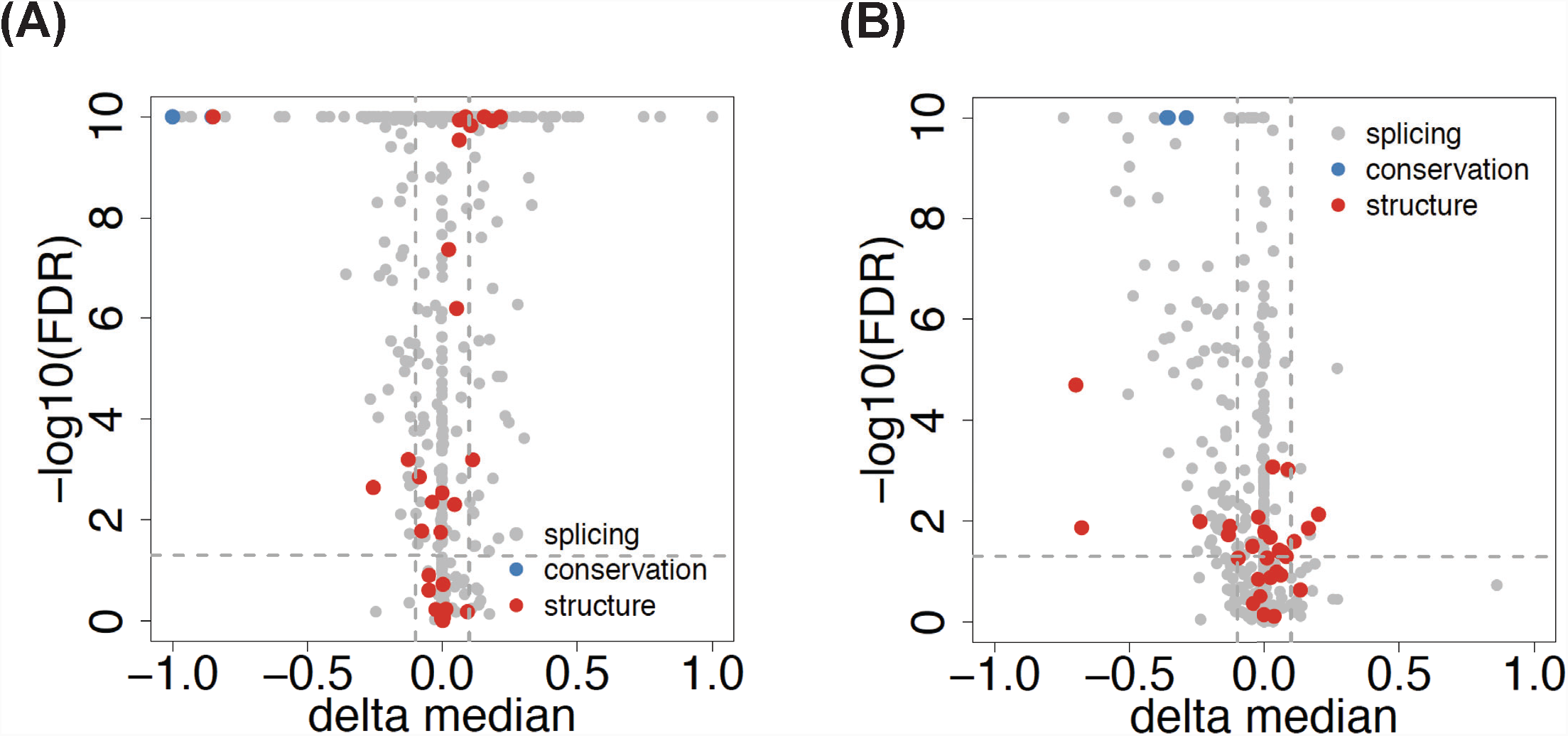
Feature evaluation. Significance of difference in feature scores between pathogenic and neutral iSNVs (**A**) on-ss iSNVs, (**B**) off-ss iSNVs. Dots represent individual features in the three categories - splicing (grey), protein structure (red), and evolutionary conservation (blue). See also Supplemental Table S1 for details. The x-axis is the median score difference, which was defined as: *median score of pathogenic iSNVs – median score of neutral iSNVs*. Feature scores were calculated as described in Methods. The y-axis is -log10 of Wilcoxon rank-sum test p-value (Benjamini-Hochberg, i.e. FDR adjusted).

### Disease-causing iSNVs affect alternative splicing

To evaluate the impact of an iSNV on splicing regulation, we considered two measures: the splice-junction scores associated with proximal exons in both 5’ and 3’ directions, as well as the deviation of the junction score of the variant from the reference allele; and the iSNV-induced difference in RBP binding affinity. The splice-junction score was computed by position weighted matrices (PWMs) measuring sequence features around the canonical junction sites (see Methods section for details) [28]. Higher scores are more likely to include the corresponding exons in the resulting mRNA. Our results showed that pathogenic iSNVs were more frequently associated with lower splice-junction scores. The median junction score of pathogenic on-ss iSNVs was 7.37 and 7.11 for donor and acceptor sites, respectively, compared to 7.95 and 7.84 for neutral iSNVs (adjusted p-value 0.017 and 1.35*10^-7^, respectively) (Table S1). For pathogenic off-ss iSNVs, the median junction scores were 7.63 and 7.95 for donor and acceptor sites, compared to 8.13 and 8.54 for neutral off-ss iSNVs (adjusted p-value 0.018 and 9.0*10^-4^, respectively) (Table S1). In addition to the junction scores, the deviation of the junction score resulting from each on-ss iSNV was calculated. The median deviation values for pathogenic iSNVs were -2.96 and -2.23 for donor and acceptor sites, respectively. The magnitude of these deviations was significantly larger than those of neutral iSNVs, which were 0.034 and -0.059 for donor and acceptor sites, respectively (Figure S3). The adjusted Wilcoxon rank-sum test p-values for donor and acceptor sites were 1.63*10^-211^ and 1.28*10^-143^, respectively (Table S1 and Figure S3). These results indicate that pathogenic iSNVs are strongly associated with exon skipping.

To evaluate the impact of an iSNV on RBP binding, we computed both the magnitude and probability of binding score changes caused by the iSNVs for 201 RBPs with known PWMs. We found that pathogenic and neutral iSNVs showed significant differences in the binding of many RBPs. Generally, the pathogenic on-ss and off-ss iSNVs induced large binding score changes in 191 and 176 RBPs, respectively, compared to the respective neutral iSNVs. Specifically, 137 RBPs showed significant differences in iSNV-induced binding scores between on-ss pathogenic and neutral variants, whereas 43 RBPs showed significant differences between off-ss pathogenic and neutral variants (adjusted Wilcoxon rank-sum test p-value < 0.05; Table S1 and Figure S4). Taken together, these findings provide strong evidence for pathogenic iSNVs affecting splicing regulation resulting in altered mRNA structures.

### Disease-causing iSNVs are associated with exons encoding functionally important protein domains

To evaluate the impact of iSNVs on protein function, we examined the structural features corresponding to potential alternatively spliced exons. We hypothesized that pathogenic iSNVs disrupt the splicing of exons that encode key protein structural domains. We captured the protein structural features for the closest neighboring exons of the iSNVs including intrinsic disorder score, secondary structure (e.g. alpha helix, beta sheet or random coil), and solvent accessible surface areas (ASA) (Table S1) [29, 30]. We also calculated the overlap percentage of the target exon with known protein domains, as well as the number of known post-translational modification sites within the exon-encoded protein domain (Table S1) [31, 32].

We found that exons in proximity to pathogenic iSNVs were more likely to encode protein domains that had lower average disorder scores and contained longer structured regions, compared to exons proximal to neutral iSNVs (adjusted Wilcoxon rank-sum test p-value 2.03×10^-11^ and 0.0497 for on-ss and off-ss iSNVs, respectively) (Table S1 and Figure S5). This result indicates that pathogenic iSNVs have a higher probability of being proximal to exons encoding structured regions. In addition, proximal exons to pathogenic iSNVs had significantly smaller average ASA scores (adjusted p-value 2.90×10^-13^ and 0.0103 for on-ss and off-ss iSNVs, respectively), which indicates that they are more likely to encode regions in the protein core as opposed to regions on the protein surface. Moreover, the closest exons to pathogenic iSNVs encoded a significantly higher percentage of residues that overlapped with known protein domains (adjusted p-value 1.91×10^-16^ and 1.98×10^-5^ for on-ss and off-ss iSNVs, respectively). Taken together, these results show that exons proximal to pathogenic iSNVs are more likely to encode functionally important protein regions. On the other hand, our analysis also suggests that protein structural features provide valuable information for prioritizing disease-causing iSNVs.

### Disease-causing iSNVs localize to conserved regions

Previous studies showed that evolutionary conservation is an important feature in assessing the disease-causing potential of SNVs [33, 34]. To determine whether an iSNV was evolutionarily conserved, we calculated the PhyloP 100-way conservation score of the iSNV locus, and the mean conservation score of a region flanking either side of the candidate iSNV by a length of 3 bp as well as 7 bp. Our results showed that pathogenic iSNVs had significantly higher PhyloP conservation scores than neutral iSNVs (Figure S6). The median conservation score for on-ss pathogenic iSNV loci was 3.08, which was significantly higher than the score of -0.03 for neutral on-ss iSNVs (adjusted Wilcoxon rank-sum test p-value <1×10^-300^). Likewise, the median score for off- ss pathogenic iSNV loci was 0.31, compared to -0.28 for neutral off-ss iSNVs (adjusted p-value 3.21×10^-38^). The large positive PhyloP scores for the pathogenic iSNV loci suggested that they evolved much more slowly than the neutral loci. Conservation scores for the 3-bp and 7-bp flanking regions were consistent with the iSNV loci scores. For the 3-bp flanking regions around the on-ss and off-ss pathogenic iSNVs, the respective medians of their mean conservation scores were 2.86 and 0.23, compared to 0.58 and -0.07 for the neutral iSNVs (adjusted p-values 1×10^-300^ and 5.47×10^-23^, respectively). Similar results were also found for the 7-bp flanking regions. These findings indicate that iSNVs at more conserved loci are more likely to be pathogenic.

### regSNPs-intron model building, performance, and evaluation

Based on the splicing, protein structure, and evolutionary conservation features described above (Table S1), random forest classifiers were built for on-ss and off-ss iSNVs, respectively. The models were built on the training set (2/3rd of original dataset) and their predictive powers were evaluated on the validation set (1/3rd of original dataset). Hyperparameters, such as number of trees and maximum depth, were optimized via the grid search with 3-fold cross-validation on the training set. For on-ss iSNVs, the random forest model contained 52 trees with the maximum depth of 13. For off-ss iSNVs, 59 trees with the maximum depth of 20 were built. The resulting on-ss and off-ss models constitute regSNPs-intron.

Based on the validation set (1/3rd of the original dataset, not used in model training), regSNPs-intron reached an AUROC (area under the receiver operating characteristic curve) of 0.96 and a Matthews correlation coefficient (MCC) of 0.79 for on-ss iSNVs and outperformed both SPANR (AUROC 0.77) and CADD (AUROC 0.81). For off-ss iSNVs, regSNPs-intron AUROC was 0.84 (MCC 0.52), compared to SPANR and CADD AUROCs of 0.54 and 0.69, respectively (Figure 3A and 3E). Our results suggest that inclusion of protein structural features encoded by the proximal exons, which are not used by either SPANR or CADD, significantly increases the performance of regSNPs-intron in predicting variant pathogenicity.

**Figure 3.**
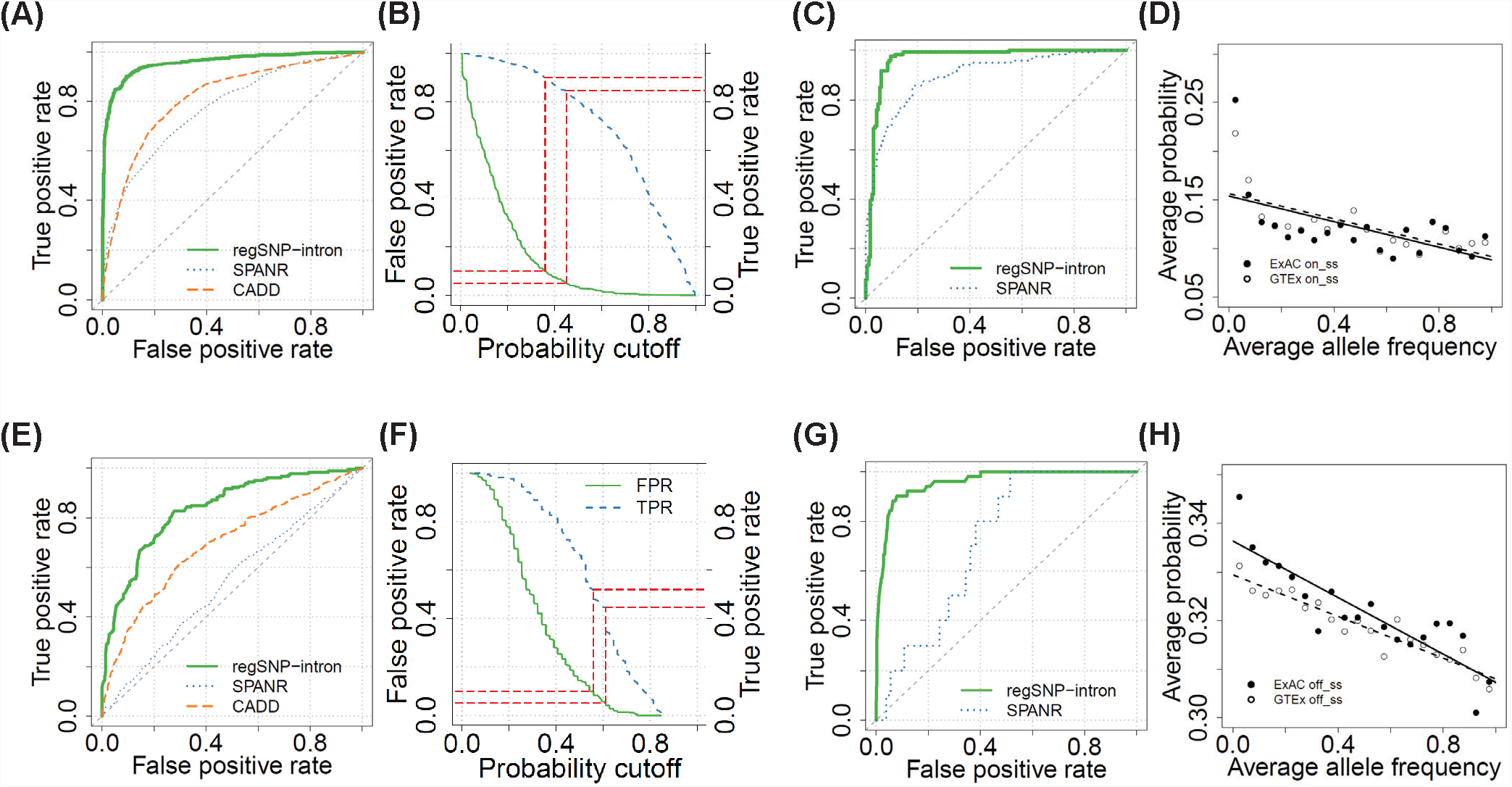
Model performance, evaluation and analysis. (**A** - **D**): On-ss iSNVs. (A) ROC of regSNPs-intron (solid green), SPANR (dotted blue) and CADD (dashed orange) on the validation set. (**B**) Selected probability cutoffs based on the false positive rate (solid green) and true positive rate (dashed blue). Dashed red lines indicate FPR = 0.05 and 0.1, respectively. iSNV with FPR < 0.05 are *Damaging*, and those with 0.05 ≤ FPR < 0.1 are *Possibly Damaging*. iSNVs with FPR ≥ 0.1 are *Benign*. (**C**) ROC of regSNPs-intron (solid green) and SPANR (dotted blue) on the independent test set of ClinVar. CADD was excluded since ClinVar was used in its model training. (**D**) Correlation between disease-causing probability and allele frequency. 20 bins were divided based on allele frequencies of the iSNVs collected from ExAC and GTEx. The x- and y-axes are the average allele frequency and predicted disease-causing probability for each bin. (**E** - **H**): Off-ss iSNVs, same as (A - D).

To further evaluate predictive powers of the features related to splicing, structure and conservation, we built separate models based on features from each of these three categories. For on-ss iSNVs, the AUROCs for splicing, structure and conservation features were 0.92, 0.72 and 0.92 respectively; for off-ss iSNVs, the AUROCs were 0.75, 0.63 and 0.68 respectively (Figure S7). These results demonstrate that each category of features provides important information in model prediction, and thus, the combination of all three categories yields the highest performance.

To control the false positive rate (FPR) of the prediction results, we reported the iSNVs with FPR < 0.05 as *Damaging*, iSNVs with 0.05 ≤ FPR < 0.1 as *Possibly Damaging* and iSNVs with FPR ≥ 0.1 as *Benign*. The reported Damaging category had true positive rates (TPR) of 0.85 and 0.45 for on-ss and off-ss iSNVs, respectively, whereas TPRs for the Possibly Damaging category were 0.90 and 0.52 for on-ss and off-ss iSNVs (Figure 3B and 3F).

### Evaluation of model performance using an independent test set

We further evaluated regSNPs-intron performance with an independent dataset from ClinVar (described above). All the ClinVar iSNVs that were also observed in HGMD or 1000 Genomes datasets were excluded from the training set to avoid overfitting. Consistent with the results described above, the regSNPs-intron model showed better performance compared to SPANR. The regSNPs-intron AUROCs were 0.96 and 0.95 for on-ss and off-ss iSNVs, respectively, whereas the SPANR AUROCs were 0.89 and 0.72 for on-ss and off-ss iSNVs, respectively (Figure 3C and 3G). We did not include CADD in this comparison since the ClinVar data were used in its original model training [18]. These results suggest that regSNPs-intron exhibits stable performance and higher prediction accuracy compared to SPANR over different datasets.

### Allele frequency was inversely correlated with disease-causing probability

Generally, allele frequency in the population should reflect the importance of the biological function of a variant [35-39]. Therefore, we examined the relationship between allele frequency and the predicted disease-causing probability of iSNVs obtained from the Exome Aggregation Consortium (ExAC) and the Genotype-Tissue Expression Project (GTEx), respectively. The iSNVs were divided into 20 bins based on their allele frequencies. For each bin, the average disease-causing probability of all the iSNVs within the bin were calculated. A strong negative correlation was observed between allele frequency and disease-causing probability for both on-ss iSNVs (ExAC R^2^ = -0.32, GTEx R^2^ = -0.47) and off-ss iSNVs (ExAC R^2^ = -0.77, GTEx R^2^ = -0.90) (Figure 3D and 3H). This result is consistent with the findings of others that variants with higher disease-causing probability are less likely to occur in the general population [20].

### Disease-causing iSNVs occur near exons associated with high disease-causing probability

We further evaluated the regSNPs-intron predictions by investigating the functional importance of exons that are proximal to iSNVs. We hypothesized that functionally important exons tolerate fewer nearby iSNVs compared to exons not associated with disease. To test this hypothesis, we extracted 75,119 exons that had at least one iSNV within ±300 bp of exon-intron boundaries based on ExAC data. One iSNV was randomly selected per exon and the disease-causing probabilities were predicted using regSNPs-intron. We observed a significant negative correlation between the predicted average disease-causing probability and the number of iSNVs near exons (R^2^ = -0.85, p-value = 6.26×10^-11^) (Figure 4). The same analysis was also performed on 160,230 exons which had at least one iSNV within ±300 bp of exon-intron boundaries based on GTEx whole genome sequencing data. A similar negative correlation was observed (R^2^ = -0.61, p-value = 0.07). This result indicates that pathogenic iSNVs tend to occur near functionally important exons.

**Figure 4.**
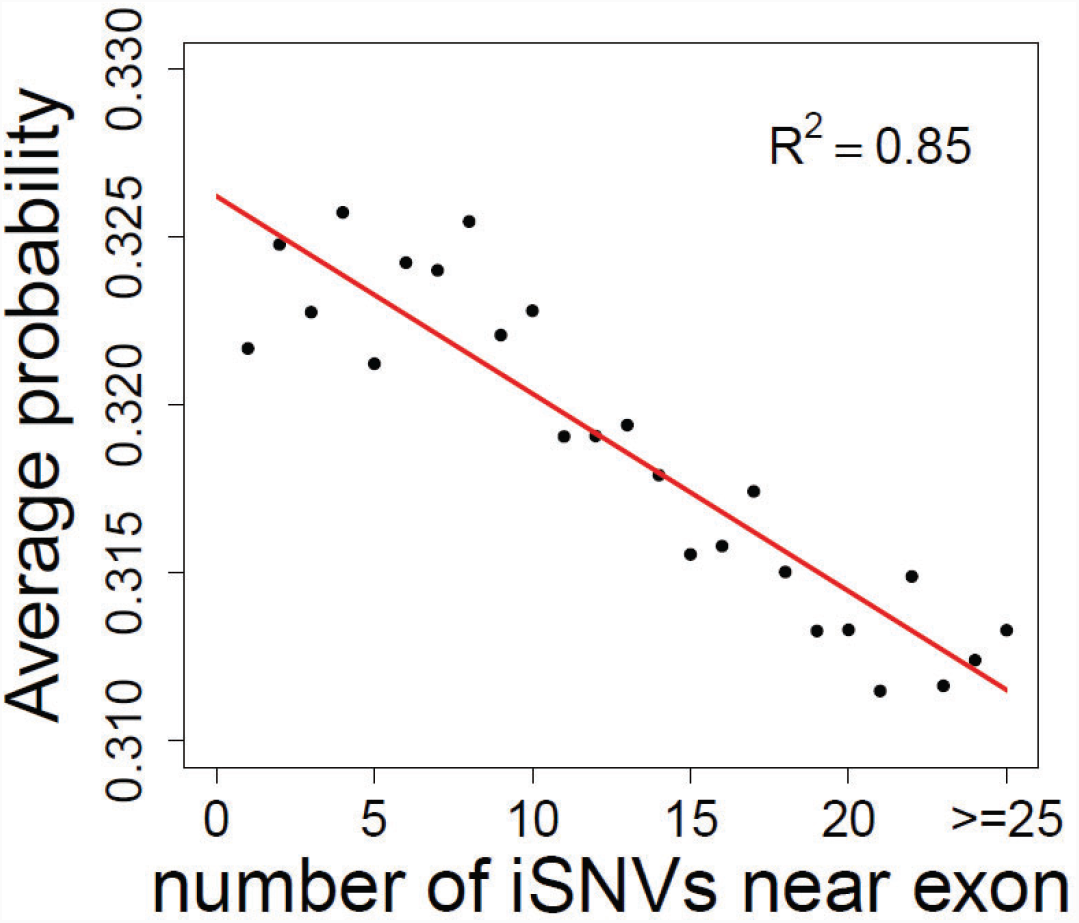
Correlation between disease-causing probability and number of nearby iSNVs for exon. The x-axis is the number of iSNV near exons; y-axis is the average predicted disease-causing probability.

### Prioritizing functional intronic variants associated with drug-induced

**cytotoxicity.** We applied regSNPs-intron to prioritize intronic variants that are associated with cellular sensitivity to clofarabine-induced cytotoxicity. We have previously performed genome-wide association studies (GWAS) for the clofarabine-response phenotype (AUC for drug-induced cytotoxicity curves) using 90 International HapMap lymphoblastoid cell lines (LCLs) of the CEU population [40]. SNVs moderately associated with clofarabine cytotoxicity (p-value ≤ 0.05) were selected as seed markers. All 17,962 iSNVs that were in linkage-disequilibrium (LD) with the seed markers and were located within 300 bp up- or down-stream of a splice junction were used in the predictions. Among these candidate variants, 622 and 84 iSNVs were predicted to be Damaging (FPR ≤ 0.05) and Possibly Damaging (0.05 < FPR ≤ 0.1), respectively (706 in total).

### Experimental validation using ASSET-seq assay

To experimentally validate the effects of the prioritized iSNVs on the splicing outcome, we designed a high-throughput functional reporter assay called ASSET-seq, that entails inserting an oligo containing an iSNV into a modified Exontrap plasmid (Figure 5A) [41]. The impact on splicing outcome of the tested iSNV was measured as the difference between the respective ratios of the sequencing reads supporting spliced and aberrant (including unspliced) transcripts for the reference and alternative alleles. The difference in the ratios for the two types of splicing outcomes between the reference and alternative alleles was analyzed by a mixed-effect model.

**Figure 5.**
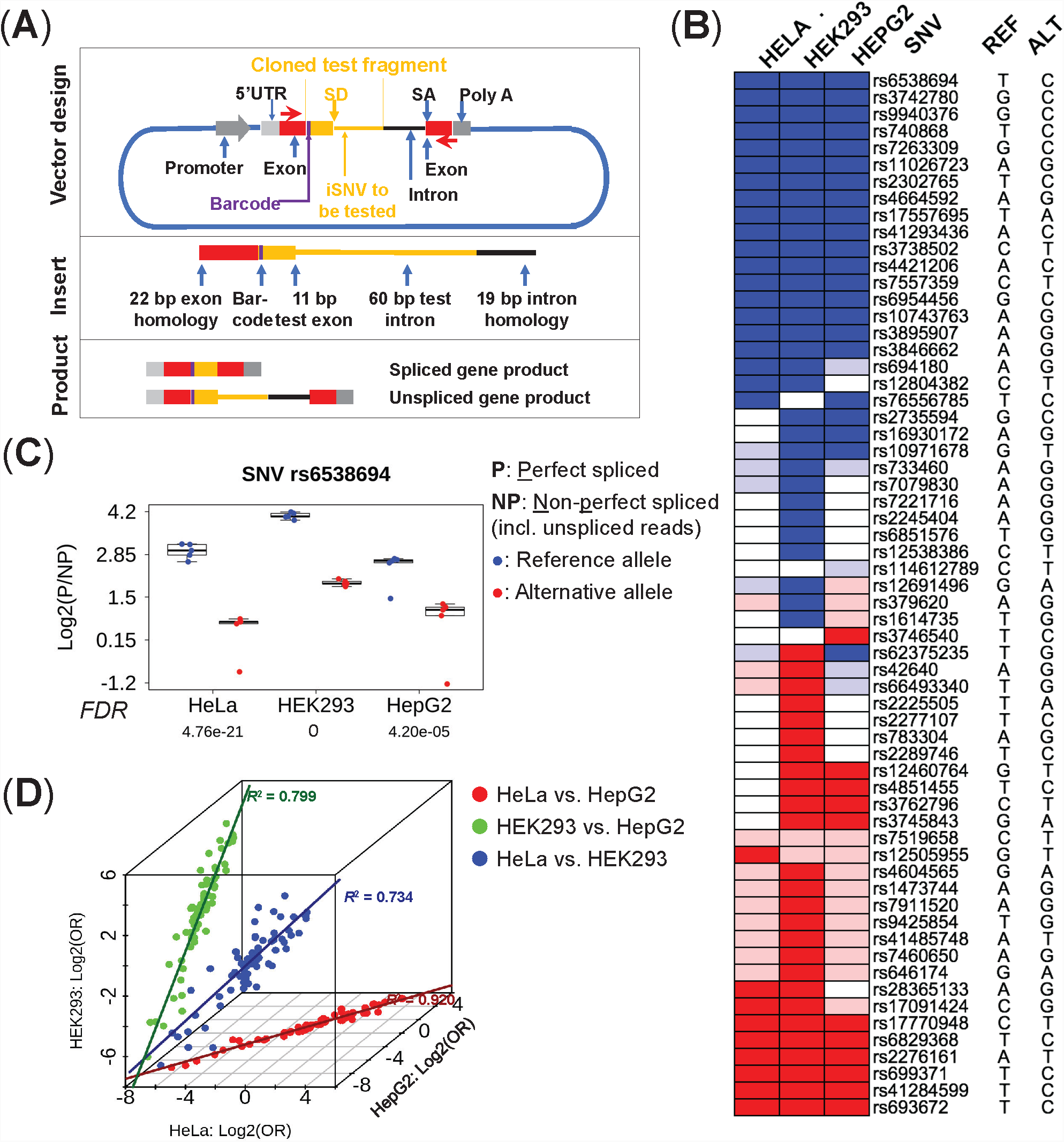
Experimental validation by ASSET-seq. (**A**) Plasmid design of the splicing assay. The insert oligo includes 11 bp of the proximal exon plus the first 60 bp of the intron containing the iSNV from individual genes (shown in orange), as well as universal plasmid exon (22 bp, red box) and intron (19 bp black line) homology segments for seamless insertion into the vector. RNA is transcribed in transfected cells from the 5’-UTR to the poly-A. PCR primers (red arrows) are used to amplify the RNA transcript. The assay produces spliced or unspliced transcripts (as well as other aberrant isoforms). Promoter: LTR RSV; SD: splice-donor site; SA: splice-acceptor site. (**B**) Summary of experimental results in three cell lines. Ref: reference allele; Alt: alternative allele. Blue color indicates an iSNV that induces a significant decrease in the spliced products (FDR ≤ 0.1); semi-blue indicates a decrease of spliced products that is not significant (FDR > 0.1). Red indicates a significant increase for spliced products whereas semi-red indicates an insignificant increase. Empty boxes are failed assays which were non-evaluable; iSNVs non-evaluable in all three cell lines (19 in total) were omitted. Validation rate (per cell line) is the percentage of assays showing a significant result in all evaluable assays. (**C**) One example of iSNV-induced alteration of splicing outcome. The iSNV (*rs6538694*) suppresses the formation of spliced gene product, as consistently indicated in all three cell lines (p-values ≤ 4.6×10^-5^). The y-axis is the log2 ratio of perfectly spliced reads (P) to aberrant (NP, non-perfect and unspliced reads). P-values were adjusted by FDR against all tested iSNVs. (**D**) Consistency of results in multiple cell lines. The x-axis is the log2 odds ratio of spliced and aberrant products (with respect to the reference and alternative alleles) for HeLa; and y- and z-axes are the log2 odds ratios for HepG2 and HEK293, respectively. Cell lines were compared in-pairwise for the iSNVs evaluable in both cell lines. Pearson’s correlations: HeLa versus HEK293 = 0.857 (blue dots, p-value = 1.02×10^-22^), HeLa versus HepG2 = 0.959 (red dots, p-value = 6.66×10^-40^), HEK versus HepG2 = 0.894 (green dots, p-value = 6.65×10^-29^). Solid lines in dark blue, red, and green are the respective regressions.

We used ASSET-seq to test the effects of 82 iSNVs on splicing outcomes in three different human cell lines: HeLa, HEK293, and HepG2. These variants were chosen from the 706 prioritized regSNPs-intron candidates (FPR ≤ 0.1) as being located in the intron on the 3′-side of the test exon and within 60 nt from the exon-intron junction, based on assay design requirements (see details in Methods). Upon removing 19 failed assays (e.g. transfection or PCR failure), the percentages of the 63 remaining iSNVs showing significant splicing impact, i.e. validation rates, for the three cell lines were: HeLa 64.4%, HEK293 96.6%, and HepG2 64.7% (FDR ≤ 0.1 in accord with the FPR; Figure 5B and Supplemental Table S2). One specific example, *rs*6538694 in the *HAL* gene, is shown in Figure 5C. Across all three cell lines (5 replicates each), a significantly higher percentage of spliced gene products was observed for the reference allele compared to the alternative allele (p-values ≤ 4.6×10^-5^). For the reference allele, the average percentage of sequencing reads supporting the spliced gene product was 88.4%, while this percentage dropped to 57.1% for the alternative allele. Moreover, we observed high consistency for the impacts of individual iSNVs on splicing outcomes across the multiple cell lines (Figure 5D).

## Discussion

The major conclusions of the current study are that RNA-splicing, protein-structural and evolutionary-conservation features all contribute to iSNV pathogenicity characterization. By integrating these three categories of feature, the regSNPs-intron algorithm efficiently evaluates the disease-causing probabilities of iSNVs *in silico*. These conclusions are based on the following evidence. First, we demonstrated that disease-causing iSNVs affect alternative splicing, localize to conserved genomic regions, and are associated with functional domain-encoding exons. Next, we provided strong evidence that regSNPs-intron has superior accuracy in computing the disease-causing probabilities for iSNVs compared to SPANR and CADD, based on 1000 Genomes and HGMD data, as well as with independent ClinVar data. Furthermore, we applied regSNPs-intron to a GWAS dataset of drug-cytotoxicity and experimentally validated the prioritized iSNVs via ASSET-seq. Taken together, this evidence strongly supports the overall concept that the regSNPs-intron algorithm, combined with the ASSET-seq assay, will facilitate studies on the regulatory functions of iSNVs and their potential roles in disease and/or drug response.

Although information on variant-induced disruption of splicing and variant conservation has been used to evaluate the impact of synonymous variants [17], our previous studies have shown that protein structural features greatly improve the prioritization of pathogenic micro-insertions/deletions as well as alternative-splicing events [20, 21]. Prompted by our earlier findings, we proposed that protein-structural features might also be informative in predicting the pathogenic effects of iSNVs. This idea was also supported by the finding that pathogenic iSNVs tend to be localized in the vicinity of exons encoding functionally important protein domains. Following the common practice of integrating multi-level features, as in algorithms such as SPANR and CADD, regSNPs-intron constitutes the first bioinformatics tool specifically designed to predict pathogenic iSNVs.

Intronic variants are typically identified by whole-genome sequencing (WGS), but they can also be captured by whole-exome sequencing (WES), particularly those iSNVs close to splice junctions that may be functionally important. To estimate the number of these intronic variants, we surveyed all the genetic variants documented in the ExAC database, which includes high-quality exome sequencing data from 60,706 unrelated individuals from a variety of large-scale sequencing projects such as the NHLBI exome sequencing project (ESP) and the 1000 Genomes project [42]. Among 7,908,659 documented SNVs in the ExAC database, these iSNVs account for 52.2% (4,126,724); therefore, identifying which of these iSNVs is important for disease pathogenesis is crucial for human genetics research. Thus, the regSNPs-intron algorithm should serve as a valuable tool for the prioritization of intronic variants, detected through WES and WGS experiments, for functional analysis.

In order to experimentally validate the impact of predicted pathogenic iSNVs on splicing regulation, we developed an innovative experimental approach, ASSET-seq. Although this assay was able to verify splicing alterations, there were a few limitations. Not surprisingly, results were influenced by technical noise from multiple sources, such as cell transfection, sequencing or PCR, that served to confound the analysis. Since these effects could not be entirely prevented, we assumed heterogeneity among the sample replicates, and applied the generalized mixed-effect model to characterize the significance of change in splicing outcomes (i.e. correlation between allele types and spliced products). Interestingly, most of the regSNPs-intron predicted candidates displayed significant impact on splicing regulation in multiple cell lines. Specifically, HEK293 cells exhibited an excellent validation rate and low noise; whereas, HeLa and HepG2 cells had lower validation rates, possibly due to larger data variabilities (high noise). Thus, our validation results confirm the effectiveness of regSNPs-intron in prioritizing iSNVs. In addition, although we tested a relatively small number of variants in this study, ASSET-seq can be extended to allow it to operate on a much larger scale.

In conclusion, we found that integrating RNA-splicing, protein-structural and evolutionary-conservation features lead to superior characterization of disease-causing iSNVs. Using regSNPs-intron and ASSET-seq in tandem enables the effective prioritization of disease-causing iSNVs. This is expected to accelerate the identification of pathogenic iSNVs, a core task of genome-wide sequencing studies.

## Methods

### Splicing features

A junction score for the closest exon boundary of each iSNV was calculated based on the position weight matrices (PWMs) derived from canonical splice sites [28]. The junction score was measured by summing the information contents of positions from -3 to +7 for donor sites, and positions from -13 to +1 for acceptor sites. In addition, for on- ss iSNVs, the change in junction score caused by allele substitution was also computed and used as a feature.

The impact of iSNVs on RBP binding affinity was measured based on the PWMs obtained from the RBPDB and cisBP-RNA databases [43, 44]. A total of 201 PWMs were collected and the magnitude and posterior probabilities of RBP binding changes were measured using methods described previously [19]. The matching score was calculated as:

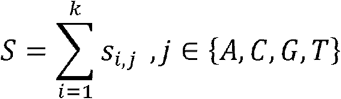

where *k* represents the number of nucleotides comprising the RBP binding site, and *s_i,j_* denotes the logarithmic ratio between the observed frequency of a specific nucleotide and random background frequency:

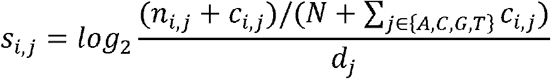

where *n_i,j_* is the count of base *j* on position *i* in the PWM, and *c_i,j_* is the pseudo-count *(√N*). *N* is the total number of binding sites used to derive the PWM, and *d*_*j*_ is the prior base frequency of base *j* (*d*_*j*_ = 0.25 for *j* = A, C, G, T).

The mean and variance of matching score distributions for binding and non-binding events were estimated as:

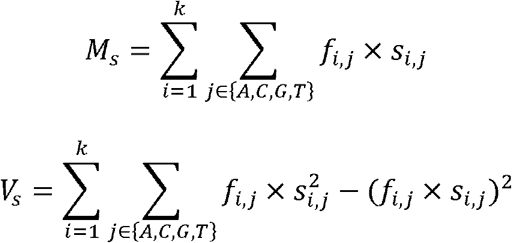

where *f_i,j_* is the approximation of the true frequency of base *j* at position *i*. For binding events 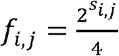, and for non-binding events *f_i,j_* = 0.25.

The magnitude of how an iSNV affects RBP binding was defined as the log-likelihood ratio between the alternative and reference alleles:

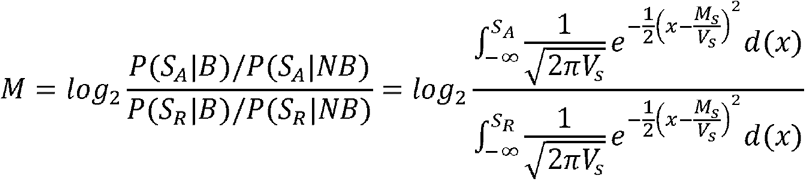

Where *S*_*R*_ and *S*_*A*_ represent the matching scores of a sequence with the reference allele and alternative allele, respectively. *P(S_A_|B)* and *P(S_R_|B)* denotes the probability of the given sequence being a binding site, while *P(S_A_|NB)* and *P(S_R_|NB)* denotes the probability of being a non-binding site.

A Bayesian-based posterior probability of the RBP binding change was calculated as:

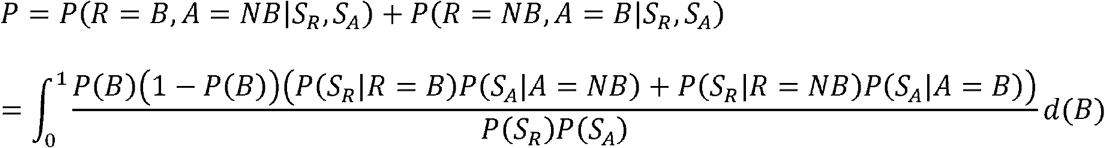

### Protein structural features

For each iSNV, protein-structural features of its closest exons were evaluated. The protein-disorder score, secondary structure and solvent accessible surface area (ASA) were precomputed for all known protein-coding genes using SPINE-D and SPINE-X [29, 30]. The known protein domains were extracted from the Pfam database [31]. Percentages of the closest exon regions that overlap with Pfam domains were measured. The post-translational modification sites (PTMs) were extracted from the dbPTM 3.0 database [32]. The number of PTM sites per 100 amino acids encoded by the closest exons were also calculated.

### Evolutionary-conservation features

Base-wise conservation scores (PhyloP) of 99 vertebrate genomes were downloaded from UCSC Genome Browser [45]. The scores on the iSNVs loci, as well as the average scores of the 3-bp and 7-bp window regions around the iSNVs, were extracted and used in machine learning.

### Machine learning model

Separate random forest classifiers were built for on-ss and off-ss iSNVs respectively. A grid search strategy with 3-fold cross-validation was used on the training set to fine-tune the hyperparameters, such as the number of trees and the maximum tree depth.

### ClinVar database iSNVs

ClinVar (version 2016/05/31) was downloaded from NCBI (ftp://ftp.ncbi.nlm.nih.gov/pub/clinvar/). We extracted the SNVs located in intronic regions. To ensure the quality of data, we only included the iSNVs that were confirmed by at least two submitters, with the exception of pathogenic off-ss iSNVs where we only required a single submitter due to the limited number of such iSNVs.

### GTEx and ExAC database iSNVs

SNVs from whole-genome sequencing data in GTEx release v6 were downloaded in the VCF format. We focused on the iSNVs within 300 bp of exon-intron boundaries. In total, there were 17,194 on-ss iSNVs and 630,557 off-ss iSNVs. Similarly, we also downloaded the variant loci from whole-exome sequencing data in ExAC r0.3.1. Corresponding allele frequencies were calculated for correlation analysis with predicted disease-causing probabilities.

### ASSET-seq plasmid construction

The modified Exontrap plasmid is shown in Figure 4A and the sequence is provided in Supplemental Text S1. The test oligos consisted of 11 bp of the upstream exon and 60 bp of the adjacent intron containing the iSNV to be tested. Additional 22-bp exonic and 19-bp intronic sequences homologous to the vector were also included for the seamless insertion of the oligos into the plasmid body. Further, a single nucleotide barcode was introduced to indicate whether the transcript came from the wild-type or variant construct. The 113-bp oligos containing different test iSNVs were synthesized in parallel as a pool using OligoMix (LC Sciences, Houston TX). In the present study, the ASSET- seq assays contained 82 pairs of reference and variant test sequences. The synthesized oligos were then cleaved from the chip, and amplified via high fidelity PCR with primers paired to the exon and intron homology sequences (Supplemental Text S1). The pooled oligos were directionally inserted into the Exontrap plasmid using the NEBuilder HiFi DNA Assembly Reaction (New England Biolabs, Ipswich MA). The assembled plasmids were transformed into bacteria and plated on LB agar plates containing ampicillin. The resulting colonies were scraped and grown in LB + ampicillin medium. Plasmid DNA was isolated using HiSpeed Plasmid Maxi kit (Qiagen, Germantown MD).

### Transfection of cell culture

The plasmid library was used to transfect three human cell lines: HeLa, HEK293, and HepG2. The cells were seeded at a density of 0.9 x 10^5^ in 24-well plates. Each plate contained 5 biological replicates per cell line. Twenty-four hours after plating, 500 ng of the library pool was transfected into each well of cells using 50 µL Opti-MEM transfection mixture containing 1.5 µL of Lipofectamine 3000 Reagent (Thermo Fisher Scientific, Waltham MA) as per the manufacturer’s instructions. Cell culture and transfection agents were used without antibiotics.

### RNA isolation and cDNA synthesis

Transfected cells were were lysed *in situ* 48 hours after transfection and total RNA was isolated using miRNeasy mini kit with the optional DNase digestion step (Qiagen, Germantown MD) following the manufacture’s protocol. Using 285-400 ng RNA, cDNA was synthesized with QuantiTect Reverse Transcription kit (Qiagen, Germantown MD) following the manufacturer’s protocol.

### Molecular barcoding

To identify the source cell line and replicate of the RNA transcripts, cDNA generated from the plasmid library in the transfected cells was PCR amplified using barcoded primers. A unique 6-nt sequence was added to the 5’-end of the forward and reverse primer for identification (Supplemental Text S1). In separate PCR reactions, 2 µL cDNA were amplified in a 50 µL volume containing 2X Invitrogen Platinum SuperFi PCR Master Mix (Thermo Fisher Scientific, Waltham MA) using 10 µM barcoded primers. PCR conditions used were 98 °C for 30 seconds, 66.6 °C for 5 seconds, and 72 °C for 15 seconds for 28 cycles. PCR samples were purified using the MinElute PCR Purification kit (Qiagen, Germantown MA) following the manufacturer’s protocol and quantified with Qubit dsDNA BR Assay kit (Thermo Fisher Scientific, Waltham MA). For sequencing by the NextSeq 500 platform (Illumina, San Diego CA), 150 ng each sample were pooled. Pooled samples also contained an equal amount of the original plasmid library with identification barcodes added by PCR as above. Pooled samples contained 20 uniquely barcoded sequence groups representing 5 biological replicates for 3 cell lines plus 5 input plasmid libraries.

### Next-generation sequencing

The pooled PCR products were sequenced using the NextSeq 500 platform (Illumina, Inc., San Diego, CA). The sequencing library was created by end-polishing the barcoded PCR products, followed by adapter ligation and amplification. The resulting library was quantified and its quality was assessed with the Agilent Bioanalyzer (Agilent Technologies, Santa Clara, CA). Approximately 90 million usable reads were generated. Raw reads were generated as fastq files for bioinformatics analysis.

### Bioinformatics analysis for sequencing data

Illumina sequencing adapters were first removed from the raw reads in the fastq files using the tool *cutadapt* (v1.9.1) [46]. Then the reads were demultiplexed into the 20 sequence groups according to the barcodes. Sequencing reads were aligned to the transcripts using STAR (<u>S</u>pliced <u>T</u>ranscripts <u>A</u>lignment to a <u>R</u>eference, v2.5.3a) [47]. The reference sequence for the alignment was built based on the plasmid (Supplemental Text S1), and read counts for the spliced and aberrant (including unspliced) transcripts were documented.

### Statistical analysis

Assuming the heterogeneity among the sample replicates, we applied the generalized linear mixed-effect model to characterize the difference of splicing patterns between reference and alternative alleles in each of the three experimental cell lines. Specifically, to determine whether there was a significant change in splicing outcome (described by counts of sequencing reads supporting the spliced and aberrant transcripts) between the two alleles, we defined *x*_*allele*_ and *x*_*splice*_ as two binary variables (i.e. 0 or 1) in the following regression equation:

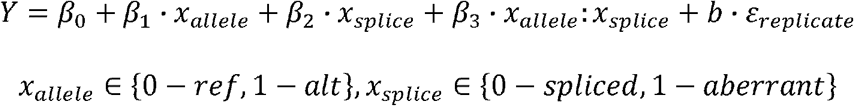

Here *Y* is the sequencing read count, which follows negative binominal distribution, and *ε* is the random effect among the multiple replicates. The impact induced by an iSNV on splicing was considered significant if the coefficient *β*_3_ was non-zero and the associated p-value was less than 0.1.

## Acknowledgements

This review was supported in part by US National Cancer Institute grant CA213466 (to YL and TS).

## Supplementary information

**Figure S1. Detailed technical protocol.**

Training data were collected from HGMD and the 1000 Genomes Project. iSNVs were first split into two classes, on-ss and off-ss respectively. 2/3 of the data were used for training, and the remaining 1/3 data were used for validation. Features were extracted and two separate random forest classifiers were built and then used to predict the disease-causing probabilities for on-ss and off-ss iSNVs. An independent test set from the ClinVar database was also used for additional model validation. The prediction performance was separately evaluated for on-ss and off-ss iSNVs. *Genomic features*: features related to DNA/RNA, e.g. exon-intron junction score, RBP binding, PhyloP sequence conservation score; *structural features*: features related to protein domains or functions, e.g. ASA score, intrinsic disorder score, secondary structure score, PTM or Pfam score.

**Figure S2. Data pre-processing.**

Apparently pathogenic iSNVs (HGMD, *top*) are closer to exons as compared to neutral iSNVs (1000 Genomes, *middle*). Thus, these iSNV data were imbalanced in terms of the distance from splice junction sites between on-ss and off-ss iSNVs. Since the numbers of on-ss iSNVs are comparable in the pathogenic and neutral groups, we needed to down-sample the huge number of off-ss neutral iSNVs to avoid potential bias in machine learning. Down-sampled off-ss iSNV data were shown at the *bottom*.

**Figure S3. Distribution of changes in splice-junction scores.**

Pathogenic iSNVs (red) had significantly lower junction scores than neutral iSNVs (black) at both acceptor splice sites (A) and donor splice sites (B).

**Figure S4. Average RBP binding score changes.**

RBP binding score changes for on-ss (A) and off-ss (B) iSNVs. Each dot represents one RBP. The x-axis is the average RBP binding score change induced by pathogenic iSNVs (*binding score with alternative allele* – *binding score with reference allele*). The y-axis is the average RBP binding score change induced by neutral iSNVs.

**Figure S5. Cumulative probability density of protein structural features.**

Pathogenic on-ss (red, A) and off-ss (red, D) iSNVs had lower disorder scores than the neutral iSNVs (black in A and D), indicating that pathogenic iSNVs are more likely to be located near exons encoding structured peptide regions. In addition, pathogenic iSNVs (on-ss, red in B; off- ss, red in E) tend to be located near exons with smaller average ASA scores as compared to neutral iSNVs (black in B and E). Moreover, pathogenic iSNVs (on-ss and off-ss, red in C and F respectively) are also more likely to be located in the vicinity of exons encoding protein regions that overlap with Pfam domains.

**Figure S6. Quantile-quantile plot of PhyloP conservation scores.**

Pathogenic iSNVs are at loci that have significantly higher PhyloP scores compared to neutral iSNVs (on-ss, A; off-ss, B).

**Figure S7. Performance of sub-models with features from each individual category.** ROCs of models trained with features only from the splicing (green), protein structural (orange), and evolutionary conservation (blue) categories respectively. (A) on-ss iSNVs, (B) off-ss iSNVs.

**Figure S8. Consistency of result data for individual iSNVs across multiple cell lines.** Consistency of measured effects for individual iSNVs on splicing outcomes are shown here. Cell lines were compared in-pairwise in terms of the log2 odds ratios of the reads supporting spliced and aberrant products with respect to the reference and alternative alleles, for iSNVs evaluable in both cell lines. Data from HeLa, HEK, and HepG2 exhibited high correlations. Specifically, the Pearson’s correlation coefficients are: HeLa versus HEK = 0.857 (blue, p-value = 1.02×10^-22^), HeLa versus HepG2 = 0.959 (red, p-value = 6.66×10^-40^), HEK versus HepG2 = 0.894 (green, p- value = 6.65×10^-29^). Solid lines in dark blue, red, and green represent regressions.

## Supplementary Tables

Table S1. Complete list of model features and their individual predictive powers.

Table S2. Additional data for ASSET-seq experiment result.

Text S1. Sequence of the modified Exontrap plasmid in ASSET-seq.

## References

1. Genomes Project Consortium, Auton A, Brooks LD, Durbin RM, Garrison EP, Kang HM, Korbel JO, Marchini JL, McCarthy S, McVean GA et al: A global reference for human genetic variation. Nature 2015, 526(7571):68–74.

2. Pagani F, Baralle FE: Genomic variants in exons and introns: identifying the splicing spoilers. Nat Rev Genet 2004, 5(5):389–396.

3. Law AJ, Kleinman JE, Weinberger DR, Weickert CS: Disease-associated intronic variants in the ErbB4 gene are related to altered ErbB4 splice-variant expression in the brain in schizophrenia. Hum Mol Genet 2007, 16(2):129–141.

4. Scotti MM, Swanson MS: RNA mis-splicing in disease. Nat Rev Genet 2016, 17(1):19–32.

5. Douglas AG, Wood MJ: RNA splicing: disease and therapy. Brief Funct Genomics 2011, 10(3):151–164.

6. Stenson PD, Mort M, Ball EV, Shaw K, Phillips A, Cooper DN: The Human Gene Mutation Database: building a comprehensive mutation repository for clinical and molecular genetics, diagnostic testing and personalized genomic medicine. Hum Genet 2014, 133(1):1–9.

7. Slaugenhaupt SA, Blumenfeld A, Gill SP, Leyne M, Mull J, Cuajungco MP, Liebert CB, Chadwick B, Idelson M, Reznik L et al: Tissue-specific expression of a splicing mutation in the IKBKAP gene causes familial dysautonomia. Am J Hum Genet 2001, 68(3):598–605.

8. Cheishvili D, Maayan C, Smith Y, Ast G, Razin A: IKAP/hELP1 deficiency in the cerebrum of familial dysautonomia patients results in down regulation of genes involved in oligodendrocyte differentiation and in myelination. Hum Mol Genet 2007, 16(17):2097–2104.

9. Neklason DW, Solomon CH, Dalton AL, Kuwada SK, Burt RW: Intron 4 mutation in APC gene results in splice defect and attenuated FAP phenotype. Fam Cancer 2004, 3(1):35–40.

10. Tazi J, Bakkour N, Stamm S: Alternative splicing and disease. Biochim Biophys Acta 2009, 1792(1):14–26.

11. Kashima T, Rao N, Manley JL: An intronic element contributes to splicing repression in spinal muscular atrophy. Proc Natl Acad Sci U S A 2007, 104(9):3426–3431.

12. Santoro A, Cannella S, Trizzino A, Bruno G, De Fusco C, Notarangelo LD, Pende D, Griffiths GM, Arico M: Mutations affecting mRNA splicing are the most common molecular defect in patients with familial hemophagocytic lymphohistiocytosis type 3. Haematologica 2008, 93(7):1086–1090.

13. Faustino NA, Cooper TA: Pre-mRNA splicing and human disease. Genes Dev 2003, 17(4):419–437.

14. Cogan JD, Phillips JA, 3rd, Schenkman SS, Milner RD, Sakati N: Familial growth hormone deficiency: a model of dominant and recessive mutations affecting a monomeric protein. J Clin Endocrinol Metab 1994, 79(5):1261–1265.

15. Cogan JD, Prince MA, Lekhakula S, Bundey S, Futrakul A, McCarthy EM, Phillips JA, 3rd: A novel mechanism of aberrant pre-mRNA splicing in humans. Hum Mol Genet 1997, 6(6):909–912.

16. Caciotti A, Tonin R, Mort M, Cooper DN, Gasperini S, Rigoldi M, Parini R, Deodato F, Taurisano R, Sibilio M et al: Mis-splicing of the GALNS gene resulting from deep intronic mutations as a cause of Morquio a disease. BMC Med Genet 2018, 19(1):183.

17. Xiong HY, Alipanahi B, Lee LJ, Bretschneider H, Merico D, Yuen RK, Hua Y, Gueroussov S, Najafabadi HS, Hughes TR et al: RNA splicing. The human splicing code reveals new insights into the genetic determinants of disease. Science 2015, 347(6218):1254806.

18. Kircher M, Witten DM, Jain P, O’Roak BJ, Cooper GM, Shendure J: A general framework for estimating the relative pathogenicity of human genetic variants. Nat Genet 2014, 46(3):310–315.

19. Zhang X, Lin H, Zhao H, Hao Y, Mort M, Cooper DN, Zhou Y, Liu Y: Impact of human pathogenic micro-insertions and micro-deletions on post-transcriptional regulation. Hum Mol Genet 2014, 23(11):3024–3034.

20. Zhao H, Yang Y, Lin H, Zhang X, Mort M, Cooper DN, Liu Y, Zhou Y: DDIG-in: discriminating between disease-associated and neutral non-frameshifting micro- indels. Genome Biol 2013, 14(3):R23.

21. Li M, Feng W, Zhang X, Yang Y, Wang K, Mort M, Cooper DN, Wang Y, Zhou Y, Liu Y: ExonImpact: Prioritizing Pathogenic Alternative Splicing Events. Hum Mutat 2017, 38(1):16–24.

22. Consortium GT, Aguet F, Brown AA, Castel SE, Davis JR, He Y, Jo B, Mohammadi P, Park Y, Parsana P et al: Genetic effects on gene expression across human tissues. Nature 2017, 550:204.

23. Livingstone M, Folkman L, Yang Y, Zhang P, Mort M, Cooper DN, Liu Y, Stantic B, Zhou Y: Investigating DNA-, RNA-, and protein-based features as a means to discriminate pathogenic synonymous variants. Hum Mutat 2017, 38(10):1336–1347.

24. David CJ, Manley JL: Alternative pre-mRNA splicing regulation in cancer: pathways and programs unhinged. Genes Dev 2010, 24(21):2343–2364.

25. Genomes Project Consortium, Abecasis GR, Altshuler D, Auton A, Brooks LD, Durbin RM, Gibbs RA, Hurles ME, McVean GA: A map of human genome variation from population-scale sequencing. Nature 2010, 467(7319):1061–1073.

26. Breiman L: Random forests. Mach Learn 2001, 45(1):5–32.

27. Landrum MJ, Lee JM, Riley GR, Jang W, Rubinstein WS, Church DM, Maglott DR: ClinVar: public archive of relationships among sequence variation and human phenotype. Nucleic Acids Res 2014, 42(Database issue):D980–985.

28. Itoh H, Washio T, Tomita M: Computational comparative analyses of alternative splicing regulation using full-length cDNA of various eukaryotes. RNA 2004, 10(7):1005–1018.

29. Faraggi E, Yang Y, Zhang S, Zhou Y: Predicting continuous local structure and the effect of its substitution for secondary structure in fragment-free protein structure prediction. Structure 2009, 17(11):1515–1527.

30. Zhang T, Faraggi E, Xue B, Dunker AK, Uversky VN, Zhou Y: SPINE-D: accurate prediction of short and long disordered regions by a single neural-network based method. J Biomol Struct Dyn 2012, 29(4):799–813.

31. Finn RD, Bateman A, Clements J, Coggill P, Eberhardt RY, Eddy SR, Heger A, Hetherington K, Holm L, Mistry J et al: Pfam: the protein families database. Nucleic Acids Res 2014, 42(Database issue):D222–230.

32. Lu CT, Huang KY, Su MG, Lee TY, Bretana NA, Chang WC, Chen YJ, Chen YJ, Huang HD: DbPTM 3.0: an informative resource for investigating substrate site specificity and functional association of protein post-translational modifications. Nucleic Acids Res 2013, 41(Database issue):D295–305.

33. Veltman JA, Brunner HG: De novo mutations in human genetic disease. Nat Rev Genet 2012, 13(8):565–575.

34. Liu X, Wu C, Li C, Boerwinkle E: dbNSFP v3.0: A One-Stop Database of Functional Predictions and Annotations for Human Nonsynonymous and Splice-Site SNVs. Hum Mutat 2016, 37(3):235–241.

35. Lupski JR, Belmont JW, Boerwinkle E, Gibbs RA: Clan genomics and the complex architecture of human disease. Cell 2011, 147(1):32–43.

36. Tennessen JA, Bigham AW, O’Connor TD, Fu W, Kenny EE, Gravel S, McGee S, Do R, Liu X, Jun G et al: Evolution and functional impact of rare coding variation from deep sequencing of human exomes. Science 2012, 337(6090):64–69.

37. Gorlov IP, Gorlova OY, Frazier ML, Spitz MR, Amos CI: Evolutionary evidence of the effect of rare variants on disease etiology. Clin Genet 2011, 79(3):199–206.

38. Marth GT, Yu F, Indap AR, Garimella K, Gravel S, Leong WF, Tyler-Smith C, Bainbridge M, Blackwell T, Zheng-Bradley X et al: The functional spectrum of low-frequency coding variation. Genome Biol 2011, 12(9):R84.

39. Subramanian S: Quantifying harmful mutations in human populations. Eur J Hum Genet 2012, 20(12):1320–1322.

40. Eadon MT, Wheeler HE, Stark AL, Zhang X, Moen EL, Delaney SM, Im HK, Cunningham PN, Zhang W, Dolan ME: Genetic and epigenetic variants contributing to clofarabine cytotoxicity. Human Molecular Genetics 2013, 22(19):4007–4020.

41. Kishore S, Khanna A, Stamm S: Rapid generation of splicing reporters with pSpliceExpress. Gene 2008, 427(1):104–110.

42. ExAC project pins down rare gene variants. Nature 2016, 536(7616):249.

43. Cook KB, Kazan H, Zuberi K, Morris Q, Hughes TR: RBPDB: a database of RNA-binding specificities. Nucleic Acids Res 2011, 39(Database issue):D301–308.

44. Ray D, Kazan H, Cook KB, Weirauch MT, Najafabadi HS, Li X, Gueroussov S, Albu M, Zheng H, Yang A et al: A compendium of RNA-binding motifs for decoding gene regulation. Nature 2013, 499(7457):172–177.

45. Pollard KS, Hubisz MJ, Rosenbloom KR, Siepel A: Detection of nonneutral substitution rates on mammalian phylogenies. Genome Res 2010, 20(1):110–121.

46. Martin M: Cutadapt removes adapter sequences from high-throughput sequencing reads. EMBnetjournal; Vol 17, No 1: Next Generation Sequencing Data AnalysisDO - 1014806/ej171200 2011.

47. Dobin A, Davis CA, Schlesinger F, Drenkow J, Zaleski C, Jha S, Batut P, Chaisson M, Gingeras TR: STAR: ultrafast universal RNA-seq aligner. Bioinformatics 2013, 29(1):15–21.

