## Supplementary figures and images for "RegSNPs-Intron: A computational framework for prioritizing Intronic Single Nucleotide Variants in Human Genetic Disease"

### Supplemental Figure 1

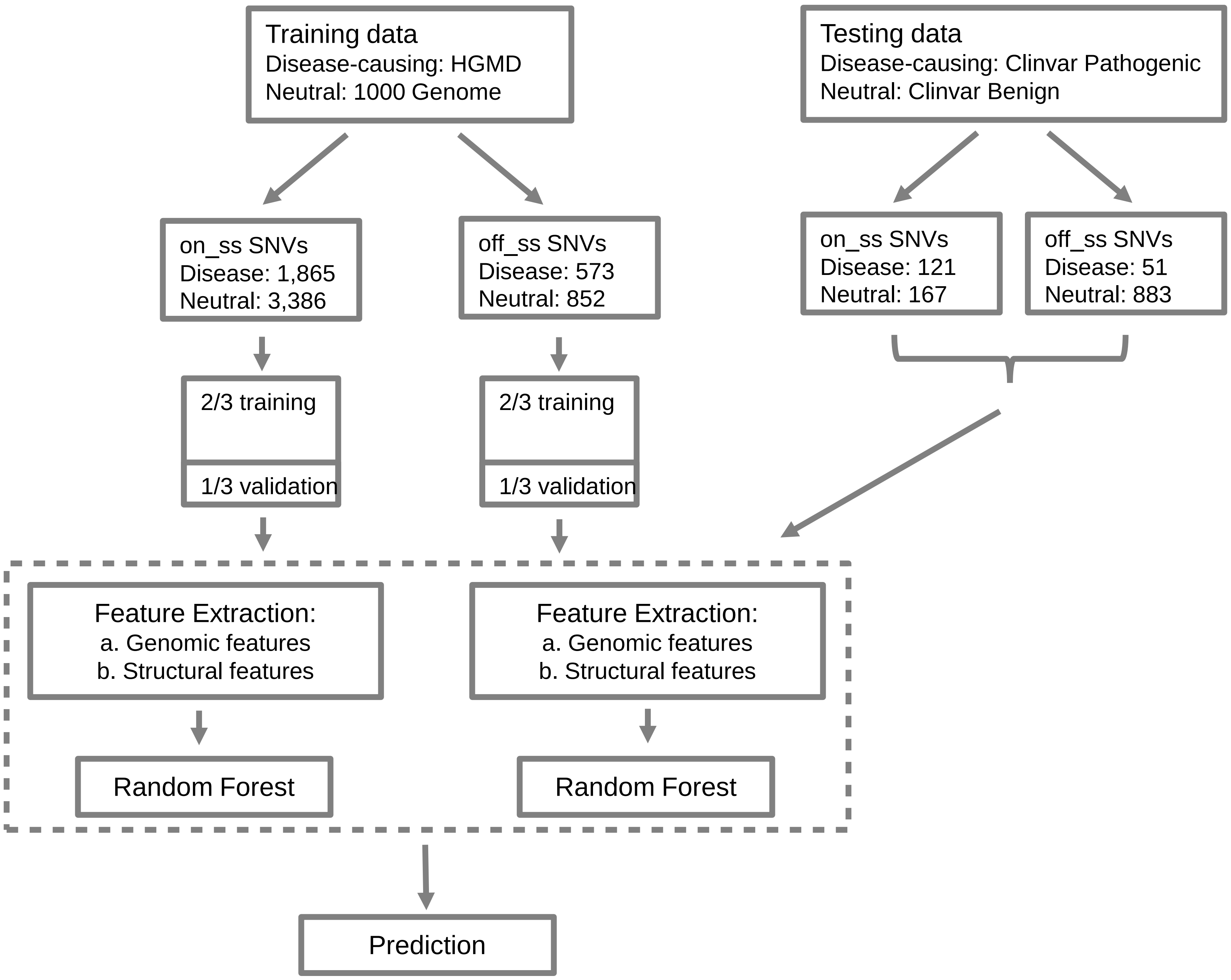

### Supplemental Figure 2

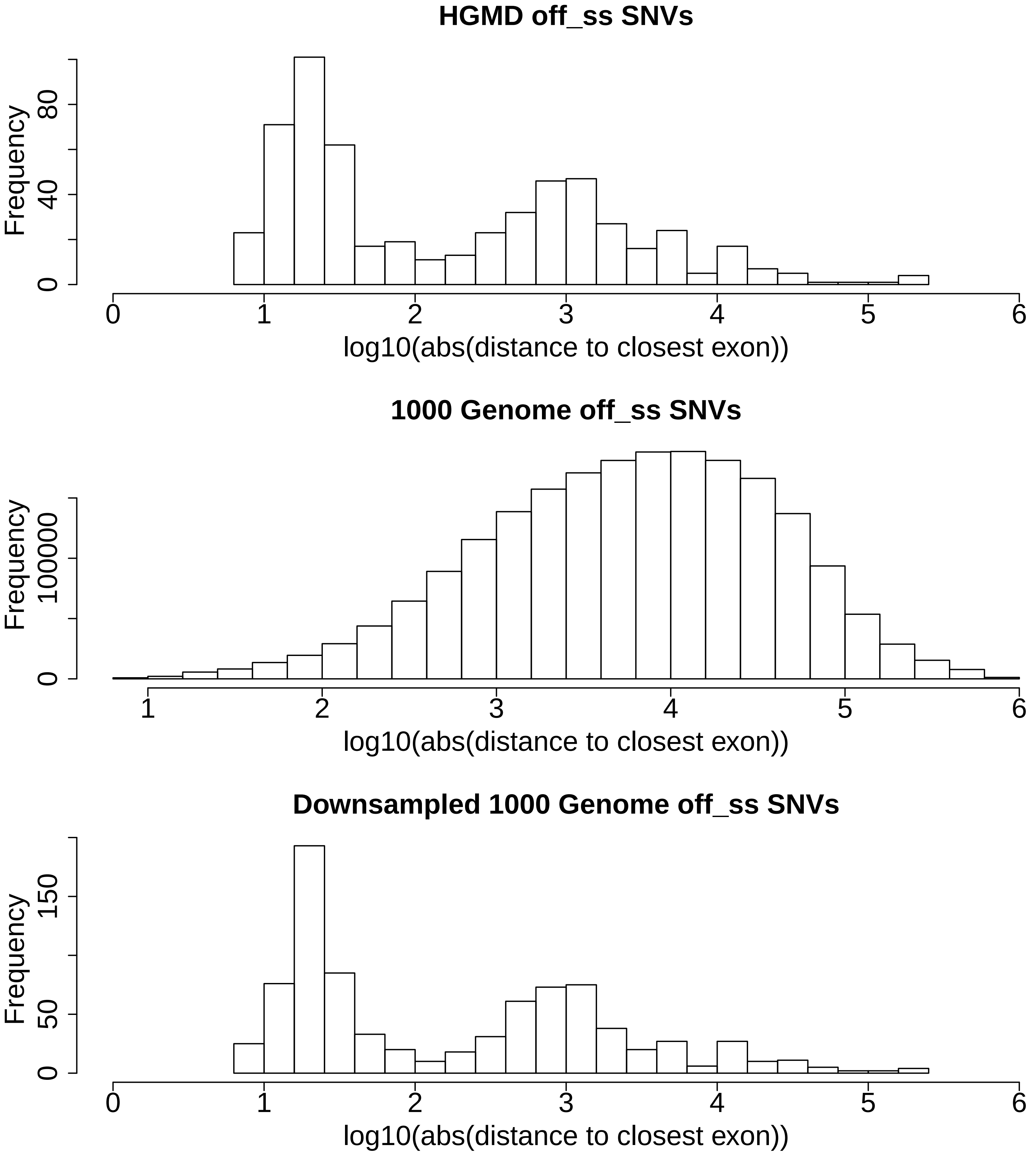

### Supplemental Figure 3

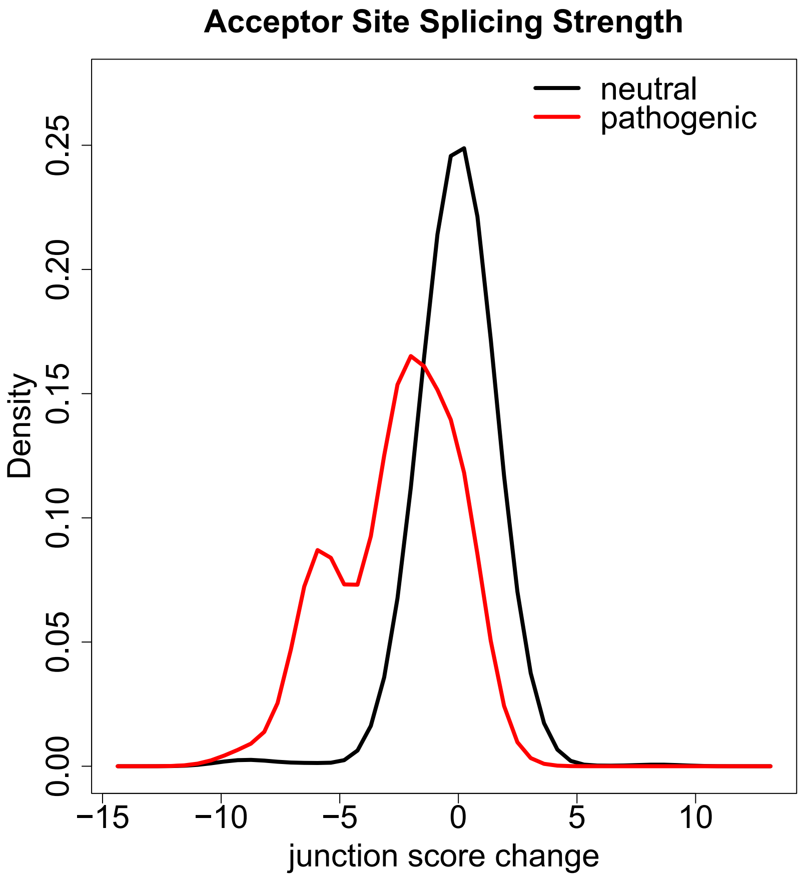

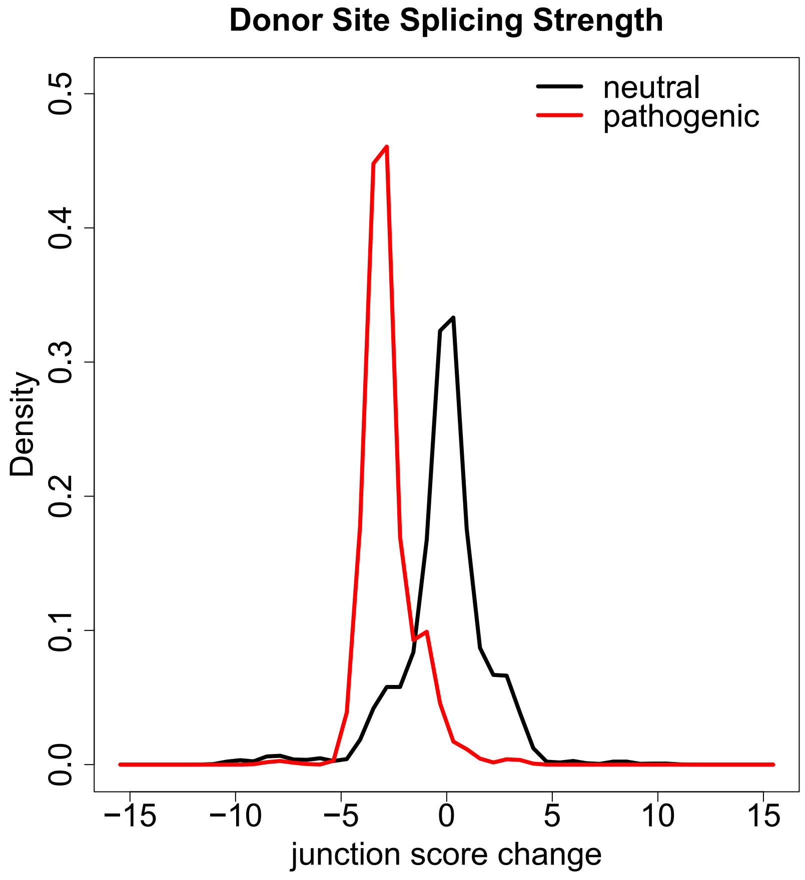

### Supplemental Figure 4

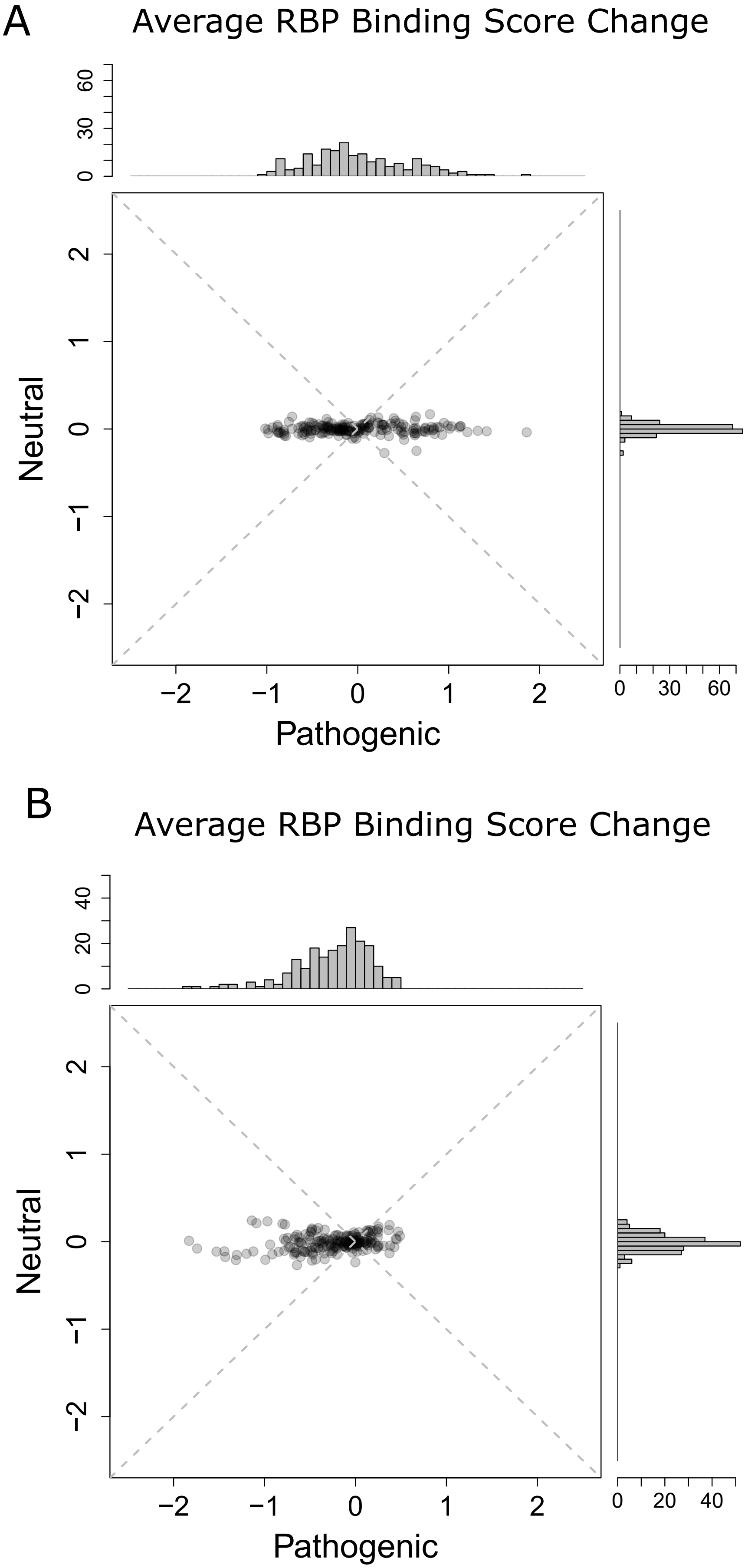

### Supplemental Figure 5

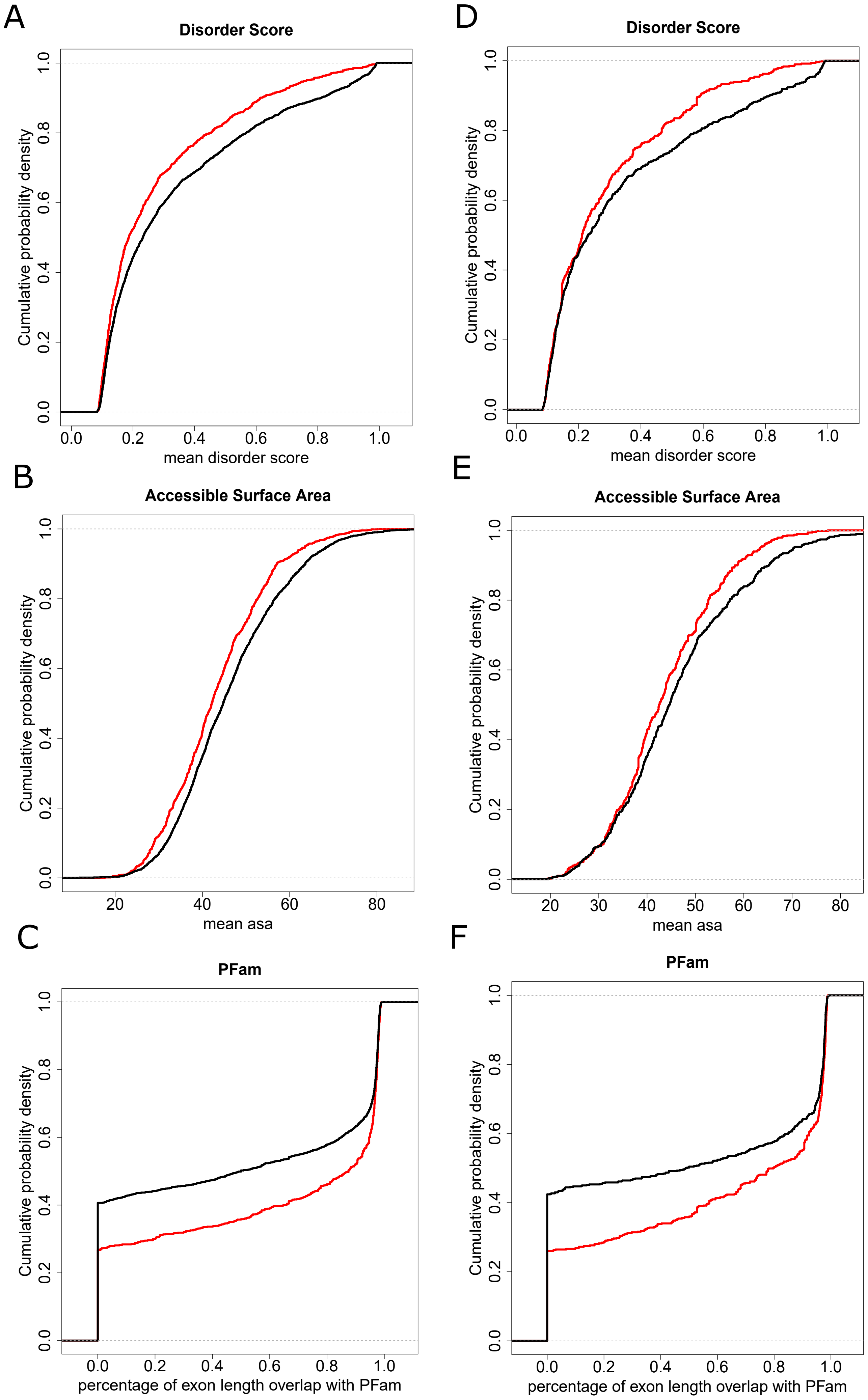

### Supplemental Figure 6

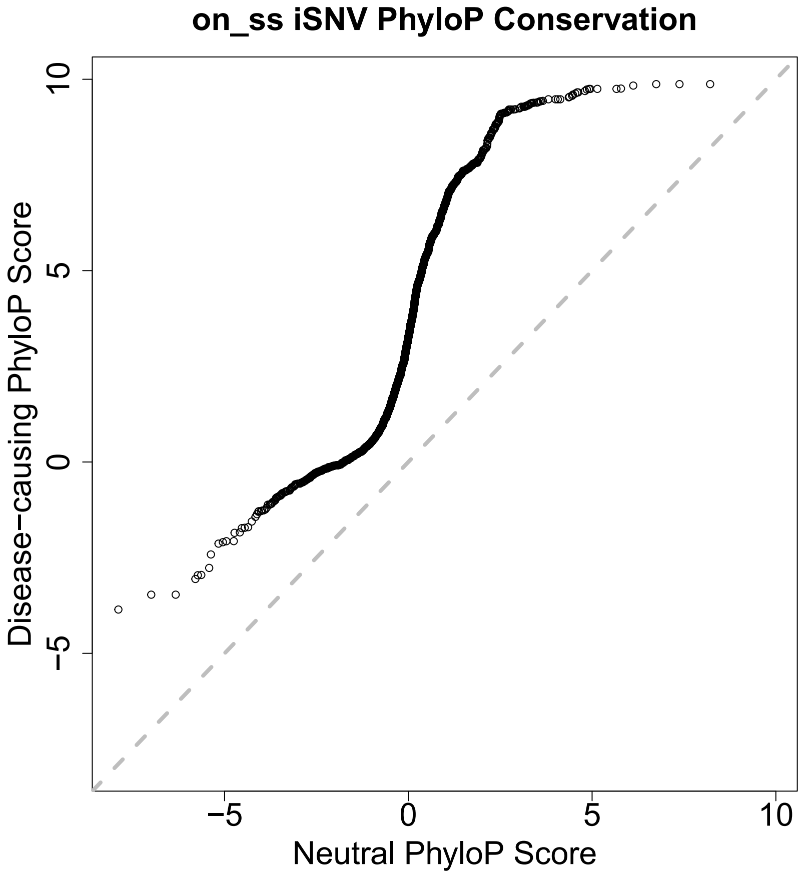

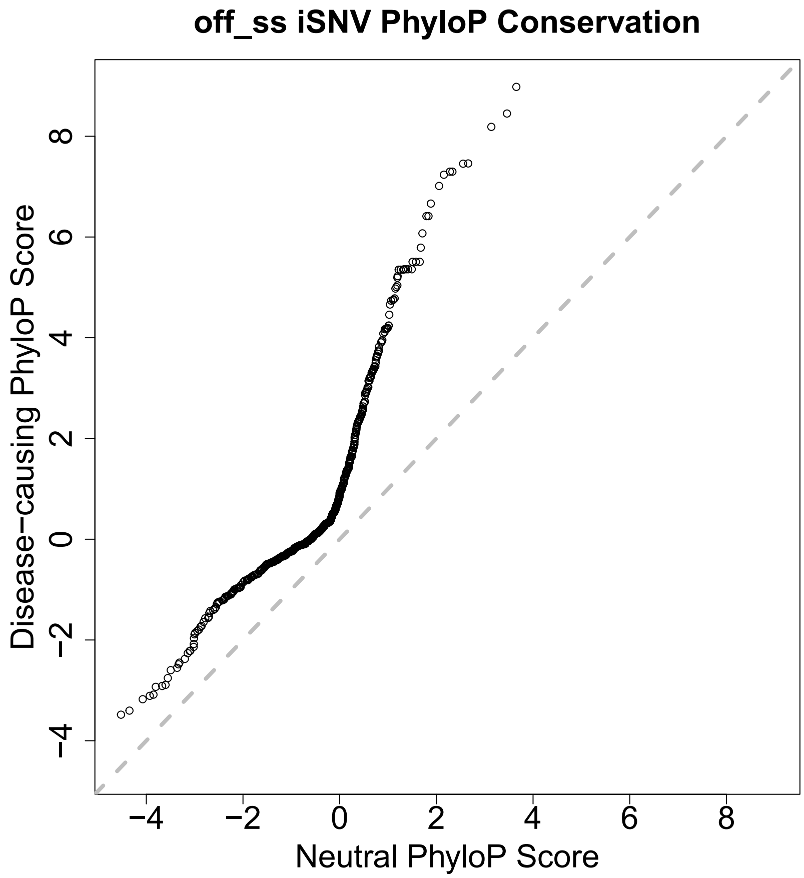

### Supplemental Figure 7

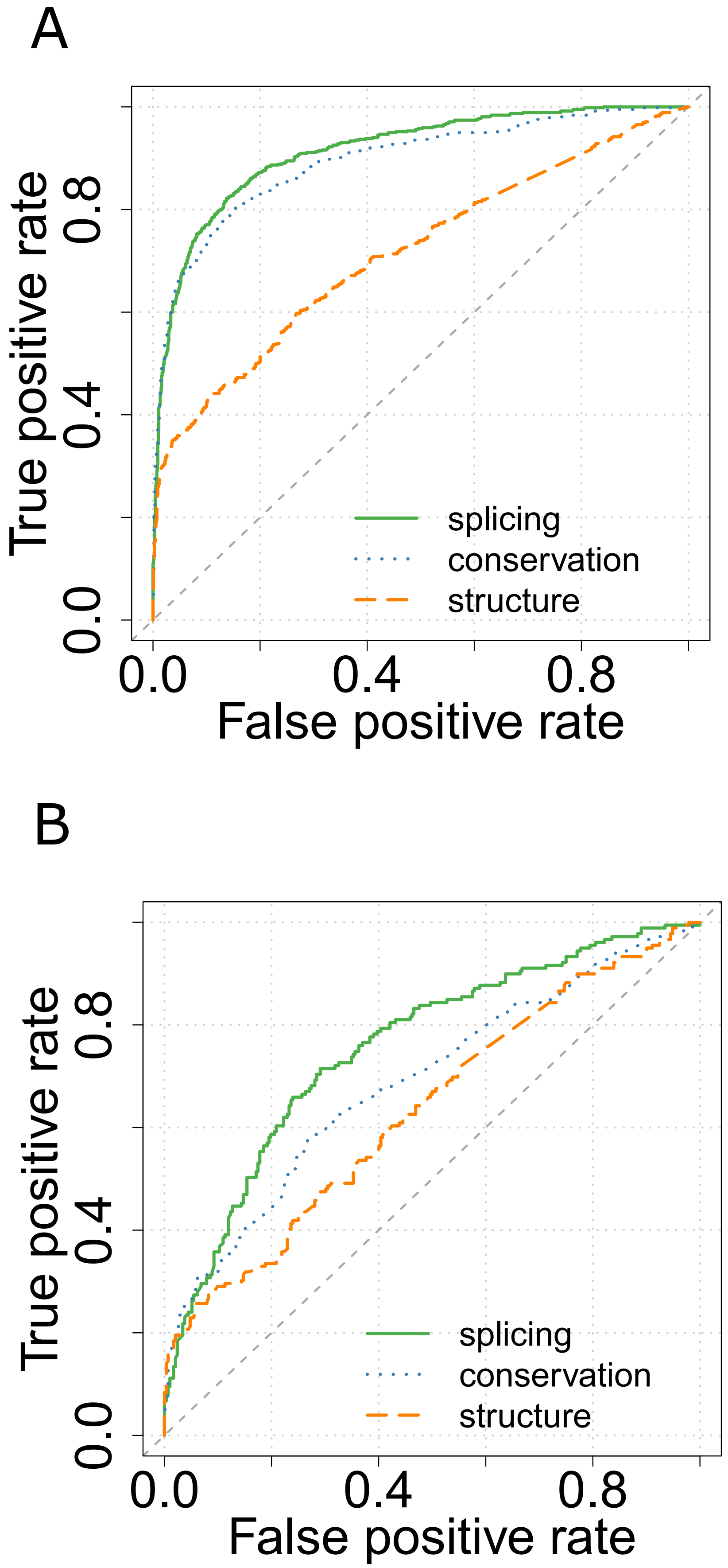

### Supplemental Figure 8

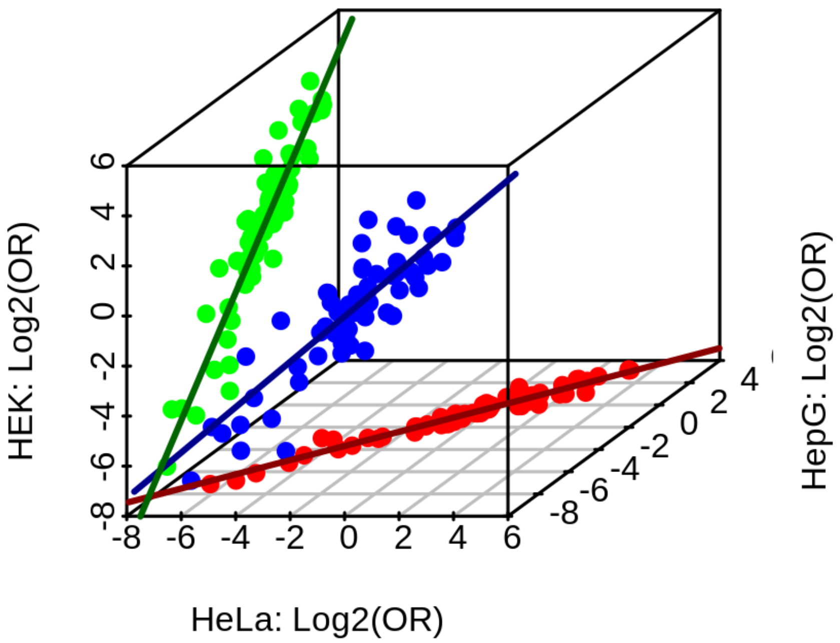
