## Supplemental Information for "RegSNPs-Intron: A computational framework for prioritizing Intronic Single Nucleotide Variants in Human Genetic Disease"

**Supplemental Text S1**

**Sequence of modified Exontrap plasmid (see also Figure 4A for graphic view)**

**KEY:**

BARCODE (FWD)

common region 1

exon homology (exon 1)

**1BC (1-nt barcode: T - ref, A - alt)**

TEST EXON **SNP** TEST INTRON (unique for every test SNP)

intron homology

common region 2

exon homology 2 (exon 2)

BARCODE (REV)

**Plasmid sequence:**

XXXXXXgcacctttgtggttctcacttggtggaagctctctacctggtgtgtggggagcgtggattcttctacacacccatgt **1BC** TEST EXON **SNP** TEST INTRON tggagctcggtacctatttggggaccccatagagcactgcactgactgagggatggtaacaggatgtgtaggttttggaggcccatatgtccattcatgaccagtgacttgtctcacagccatgcaacccttgcctcctgtgctgacttagcaggggataaagtgagagaaagcctgggctaatcagggggtcgctcagctcctcctaactggattgtcctatgtgtctttgcttctgtgctgctgatgctctgccctgtgctgacatgacctccctggcagtggcacaactggagctgggtggaXXXXXX

^*^Note: The *test exon* and *intron* is the unique part of the insert oligo. This is a fragment taken from the real gene hosting the SNP to be tested, including 11 bp of the closest exon on the 5’-side of the SNP and 60 bp of the intron containing the SNP.

ID barcodes for reads (FWD/REV in 5'-3' direction):

|  | **Input** | | **HeLa** | | **HEK293** | | **HepG2** | |
| --- | --- | --- | --- | --- | --- | --- | --- | --- |
| Replicate | Forward | Reverse | Forward | Reverse | Forward | Reverse | Forward | Reverse |
| **1** | AACGTC | AAACTC | ATCAAC | ATGCAC | GAAACC | GACAAT | TAGAAC | TATGCC |
| **2** | ACATGT | ACAACC | ATGTTG | CAATAC | GCACTA | GCGTTT | TCAAAG | TCCATA |
| **3** | ACCTTT | ACGGTT | CACAAG | CATCTA | GGTCTA | GTAATC | TCGATT | TCTACC |
| **4** | AGAAGG | AGACGT | CCAAAT | CGTTTC | GTAGAG | GTTAGT | TGCTAG | TGAACC |
| **5** | AGTGGA | AGTTAC | CTATGG | CTCCTT | TAACCC | TACAGA | TTCGAA | TTAACG |
